# When do bacterial mechanosensitive channels get activated?

**DOI:** 10.64898/2026.10.23.761342

**Authors:** S B Sai Smaran, Octavio Albarrán, Renata Garcés, Jeff D. Eldredge, Sergei Sukharev, Christoph F. Schmidt

## Abstract

Bacterial cell envelopes are complex multilayer structures that combine mechanical protection with a multitude of transport and sensory functions. Bacteria maintain substantial turgor pressures which is needed to withstand external forces, grow and maintain shape. An osmotic downshock can create fatal pressure levels. To survive, bacteria rapidly release osmolytes, mainly ions and small metabolites, through mechanosensitive channels (Mscs). Mscs are embedded in the plasma membrane (PM) and act as pressure/tension-activated valves, reducing pressure and preserving cell integrity. It is not well understood, however, how overall stresses are distributed among the layers of the cell envelope, and, in particular, when tension would be borne by the PM and felt by the channels. In this study, we numerically and analytically examine stress distributions within a two-layer model of the cell envelope under varying turgor pressure. We use experimentally determined parameters for PM and peptidoglycan (PG) elasticities, and include enzyme/antibiotic induced weakening of the PG. We introduce, as a critical parameter, a PM reservoir, without which an observed substantial area increase during the shock would not be possible without PM rupture. We used finite-element simulations to model cell swelling assuming linear elasticity of the envelope components, and analytical calculations to model nonlinear PG strain hardening. We analyzed tension sharing between PG and PM with varying PM reservoir sizes during osmotic downshocks. We estimate that an envelope area increase of slightly more than 50% would occur at a turgor pressure of 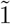1 atm and would be sufficient to fully unfold a 50% PM reservoir and also generate sufficient PM tension to activate mechanosensitive channels. A softening of the PG layer with lysozyme or *β*-lactam antibiotics is predicted to increase cell expansion at a given turgor pressure and trigger earlier activation of mechanosensitive channels, leading to increased, deleterious metabolite loss.

## 1 Introduction

The bacterial cell envelope performs a multitude of vital functions. It provides mechanical stability during cell growth and division, under varying environmental conditions. At the same time, it carries receptors, channels, and transporters that enable transport of material and signaling [1]. Bacteria precisely regulate osmolyte, dominantly ion, fluxes in order to maintain a turgor pressure of typically between about 1 to 10s of atmospheres, depending on the species and nutrient conditions [2, 3]. This pressure, acting normal to the surface, is contained by lateral tension in the cell envelope. As an example, estimates based on the van ‘t Hoff and Young–Laplace laws show that, for a rod-shaped bacterium with a transverse radius of 0.5 µm and an intracellular osmolarity excess of 200 mOsm relative to the external medium [4], the resulting turgor pressure of approximately 5 atm would generate a membrane tension of 125 mN*/*m in the hemispherical end caps. The cylindrical part would experience a hoop tension of 250 mN*/*m. This tension exceeds lytic lipid membrane tension by a factor of 30 [5–8] and therefore must be mainly borne by the other structural layers of the cell envelope. The sharing of tension between the lipid membrane(s) and peptidoglycan depends on the elastic properties and overall resting areas of those layers. Because all layers are constantly growing and remodeling, the regulation of insertion rates of new material determines this tension distribution. Note that the normal force generated by turgor pressure is transmitted through the PM to the PG, while PM and PG are sharing lateral tension.

The envelopes of Gram-positive bacteria comprise the PM and a relatively thick (20–40 nm) PG layer, separated by a polyelectrolye-filled periplasmic layer, whereas those of Gram-negative bacteria consist of the PM, a thin PG layer (2.5–4 nm), also separated by a periplasmic layer, and an outer lipid/lipopolysaccharide membrane (OM) [1, 9]. The material properties of PM and PG greatly differ. The PG layer is a covalently linked elastic network, while the PM is an incompressible 2D fluid that cannot withstand high tension without rupturing [5, 6]. It is therefore generally thought that under unperturbed, resting conditions, most of the tensile load in the cell envelope is borne by the elastic PG layer and, in Gram-negative bacteria, possibly the OM [10], shielding the PM from turgor-generated tension. To remain shielded from tensile stress while transmitting normal force to the PG, the PM must have an area greater than that of the PG. Because PG is elastic whereas the PM is essentially inextensible, the PM must actually maintain a sufficient area resrvoir to accommodate minor changes in cell volume caused by osmotic fluctuations. Thus, growth and division require tightly coordinated maintenance and expansion of the coupled envelope layers to ensure proper partitioning of turgor-generated stress and prevent membrane damage.

Bacteria survive osmotic downshocks not only through their elastic PG layer but also with the help of mechanosensitive (MS) channels in the PM, which act as rapid osmolyte-release valves, moderating turgor buildup. It was shown in *ex-vivo* experiments that the channels can be activated by lateral membrane tension [11–13]. In *E. coli*, seven genes encoding MS channels have been identified [11, 14], of which MscS and MscL provide coordinated protection against sudden osmotic downshifts [15, 16]. MscS is a low-threshold, small-conductance channel activated at moderate, sublytic tensions of 5–7 mN*/*m [17, 18], whereas MscL is a high-threshold, large-conductance channel activated at nearly lytic tensions of 12–15 mN*/*m in model membranes [18–20].

Under steady-growth conditions, the PM is expected to experiences little tension, and therefore both channels would remain closed, minimizing proton and ion leakage and loss of essential solutes. Turgor-generated cell wall tension is, in that case, borne primarily by the PG layer. Under hypo-osmotic challenge, the elastic PG will expand while the inextensible PM will loose area reservoirs. Once these are exhausted, the PM will rapidly experience substantial tension. The pressure at which this occurs, depends on the elastic properties of the PG and the amount of excess PM area, the latter of which remains poorly constrained experimentally. Once PM tension approaches the activation range of MscS, the channel can open and release osmolytes, providing a rapid mechanism for relieving turgor before the envelope experiences catastrophic failure.

Imaging wild-type *E. coli* and mutants lacking various combinations of MS channels during osmotic downshock in a microfluidic system, with an exchange time of ∼0.8 s, showed that cells swell to ∼120% of their initial volume within ∼40 s [21]. Cells lacking MscS and MscL exhibited greater swelling and slower relaxation, indicating that these channels activate rapidly in wild-type cells to limit volume expansion. Much faster dilution experiments, using stopped-flow measurements, showed that a 600 mOsm downshift from 1200 mOsm, completed within ∼8 ms, produces an initial swelling phase lasting ∼70 ms before channel activation [15, 16, 22]. Wild-type cells can undergo nearly a twofold increase in volume without loss of viability, implying that substantial PM reservoirs are available before membrane tension rises sufficiently to activate MS channels. Under the geometric assumptions used in ref. [15], this volume increase corresponds to ∼50% cell envelope expansion.

Cells lacking the major channels MscS, MscK, and MscL can undergo ∼200% volume expansion and ultimately lyse [15]. These observations demonstrate that a substantial PM area reservoir exists and that appreciable global membrane tension develops only after this reservoir is depleted. They also illustrate the close mechanical coordination between PG and MS channels in limiting volume expansion and preventing lysis. The magnitude and kinetics of the hypotonic response depend critically on PG prestress and elasticity, as well as on the size of the PM area reservoir. In constantly growing bacteria, this raises the question of how the import of osmolytes, PG growth and membrane synthesis are coordinated to maintain turgor and sufficient PM reservoirs. In Gram-positive bacteria, MS channels have been proposed to participate in the coordination between PG expansion and membrane synthesis [23, 24].

A schematic illustrating our hypothesis for how the mechanical interplay among PG, PM, and MS channels prevents cell bursting is shown in Fig. 1. We consider three scenarios. In the first, the PM and PG have an optimal area relationship. During osmotic downshift, moderate elastic stretching of PG allows the PM to unfold and accommodate volume expansion before experiencing substantial tension. This “silent swelling” buffers moderate osmotic shocks and increased turgor without releasing intracellular metabolites. With a stronger shock, continued expansion exhausts the PM area reservoir and generates sufficient membrane tension to activate MS channels, triggering osmolyte release and water efflux that reduce both intracellular pressure and volume.

**Fig. 1.**
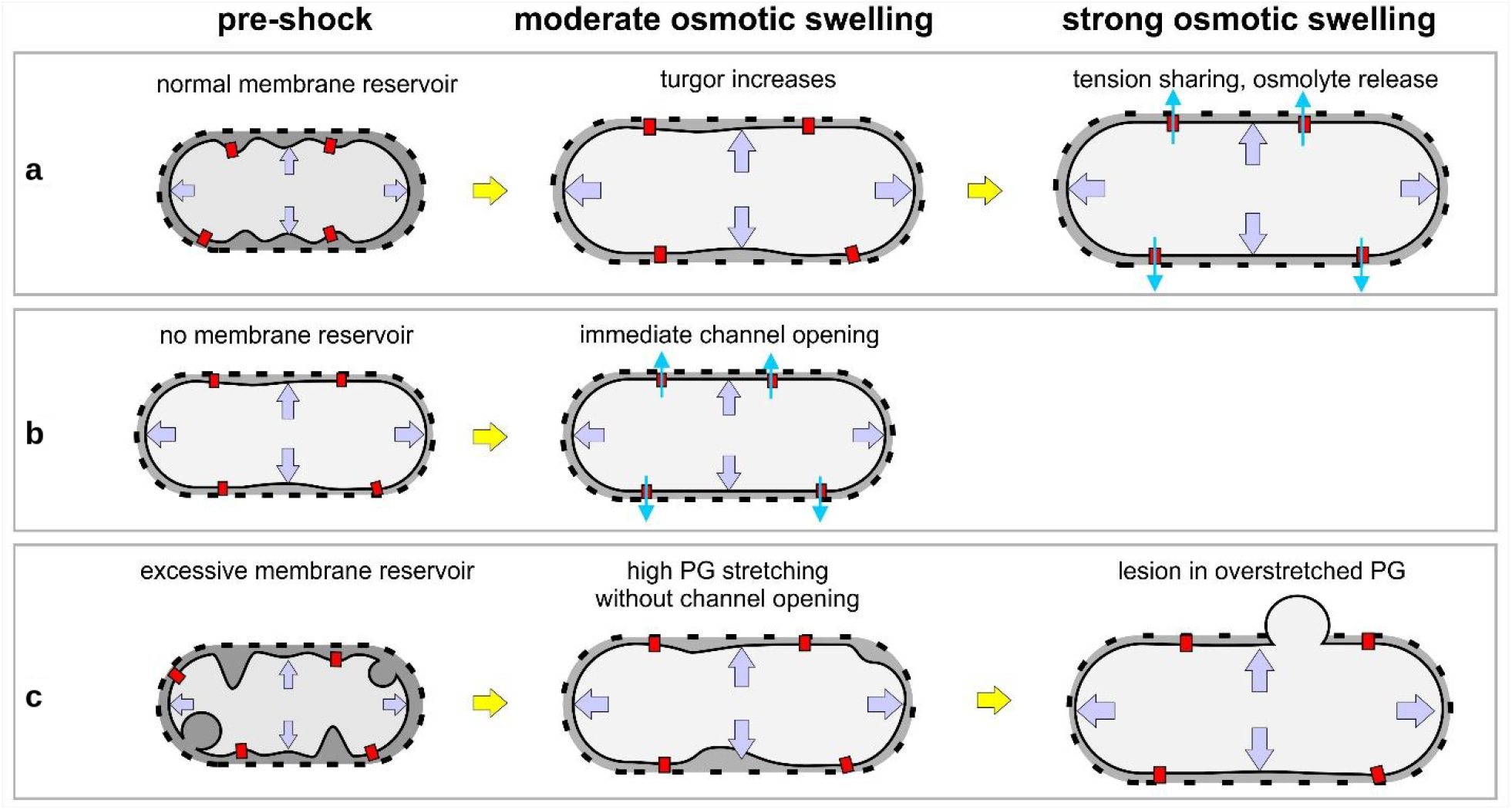
Cartoon illustrating possible scenarios of tension sharing between PG and PM under hypoosmotic shock. **a**: An optimal amount of PM excess area permits some swelling of the cell under moderate shocks without triggering MS channels, while larger shocks would trigger MS channels to prevent lytic cell swelling. **b**: In the absence of excess membrane, low sublytic shocks would lead to deleterious ion and metabolite loss. **c** A large amount of excess membrane may lead to excessive stretching of PG without triggering channel opening and protective osmolyte release, increasing the probability of lesions in PG and blebbing.

In the second scenario, the PM has little or no excess area. Water influx therefore immediately generates membrane tension, leaving less buffering capacity against elastic PG swelling. Even relatively small osmotic shocks could then activate MS channels and cause solute loss, while larger deformations could drive the PM toward failure within the PG envelope. In the third scenario, the PM has an excessive area reservoir. A larger degree of expansion would then be required to generate channel-activating tension, potentially allowing turgor to rise sufficiently to damage the PG layer and produce membrane blebbing.

The osmolarity difference between the cytoplasm and periplasm, as well as the distribution of mechanical stress among the PM, PG, and the outer membrane mechanically coupled to the PG, remain subjects of debate. Early studies suggested that the osmolarities of the periplasm and cytoplasm are nearly equal [25]. More recently, it has been proposed that the periplasm temporarily stores osmolytes released from the cytoplasm during osmotic downshock, partitioning the overall osmotic gradient between the cytoplasm and external medium and thereby buffering the osmotic pressure acting on the mechanically vulnerable PM [26].

In the present work, we simplify the system by representing the cell envelope as a two-layer structure: an osmotically active PM covered by an elastic, permeable cell wall. This representation describes the PG layer of Gram-positive bacteria and provides an effective-layer framework for the composite PG–outer membrane envelope of Gram-negative bacteria, without resolving the outer membrane explicitly, while neglecting periplasmic layers for simplicity. Using quasi-static analytical calculations and finite-element simulations with realistic material parameters, we address the fundamental question of how turgor-generated in-plane stress is partitioned between the PG and PM under resting conditions and during osmotic stress.

## 2 Methods

### 2.1 Envelope geometry and reference states

We modeled the bacterial envelope as two concentric thin spherical layers, the PM and the PG, without explicitly resolving the outer membrane or periplasmic pressure. For the illustration of fundamental principles, we neglect the actual sphero-cylindrical shape of bacteria such as *E. coli*. The calculations use PG material parameters. For simplicity, we neglect possible contributions of the OM. Because the layer thicknesses are much smaller than the cell radius, PM and PG were taken to have a common radius *r* while carrying separate tensions, *T*_PM_ and *T*_PG_. For a spherical envelope under turgor pressure *P*, the Young–Laplace law [27] gives

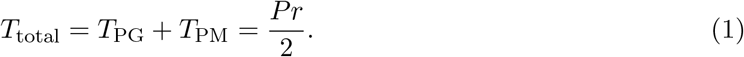

Pressure therefore determines the total wall tension, while the properties of the PG and PM determine how that tension is divided between the two layers.

We used two reference states. In the completely relaxed state, denoted by subscript *r*, there is no turgor and the wall bears no tension. With *A*_*r*_ and *r*_*r*_ the relaxed envelope area and radius, and *A* and *r* their values for a pressurized cell, area strain and radius are

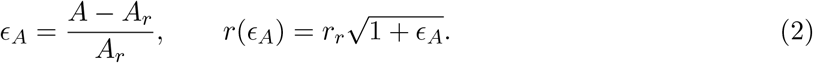

The baseline state, denoted by subscript 0, represents a Gram-negative bacterium at its resting turgor *P*_0_, with radius *r*_0_ = 0.5 µm and wall tension *T*_0_ = *P*_0_*r*_0_*/*2. We used *P*_0_ = 1 atm (*T*_0_ = 25.3 mN*/*m) for the finite-element simulations and the linear analytical model, and *P*_0_ = 0.3 atm (*T*_0_ = 7.6 mN*/*m) for the strain-hardening model. In the baseline state, the PM still retains excess area (which we will quantify with reference to this baseline state), so that all of the baseline tension is borne by the PG. The envelope area change relative to the baseline is

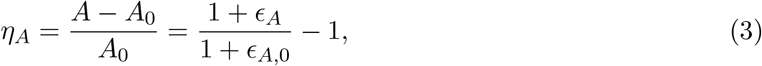

and for the spherical geometry *V/V*_0_ = (1 + *η*_*A*_)^3*/*2^.

### 2.2 Constitutive relations

In the linear model, used for the numerical simulations, the PG was treated as a linearly elastic layer with Young’s modulus *E*_PG_ = 25 MPa [28], thickness *t*_PG_ = 4 nm, and Poisson ratio *v*_PG_ = 0.3. Its tension, measured from the relaxed state, is

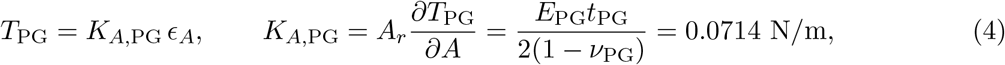

which gives a baseline area strain *ϵ*_*A*,0_ = *T*_0_*/K*_*A*,PG_ = 0.355 at 1 atm.

To include strain hardening of the PG for the analytical calculations, we replaced Eq. (4) with the nonlinear constitutive model of Albarrán et al. [29], which is based on the linear (radial) strain

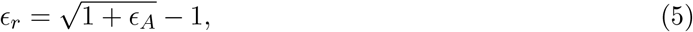

and the stiffening function

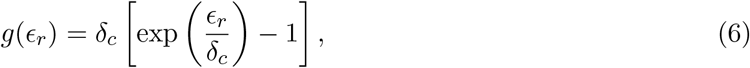

where *δ*_*c*_ sets the strain at which the response changes from linear to hardening. The PG tension becomes

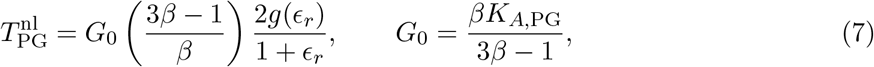

with *β* = 1 for a sphere. This choice of *G*_0_ preserves the initial slope of the linear model, so the two models differ only as the PG is extended further. We used *δ*_*c*_ = 0.08; for the baseline we solved 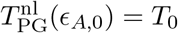, which gives *ϵ*_*A*,0_ = 0.086 at 0.3 atm, and set 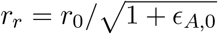.

To describe the excess area of the PM, we defined a wall area *A*_*c*_ above which the membrane is taut and bears tension, and the reservoir size as the fractional excess of *A*_*c*_ over the baseline area,

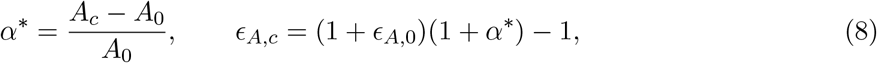

where *ϵ*_*A,c*_ is the corresponding area strain. We assume that the PM tension depends linearly on strain once the membrane reservoir is exhausted,

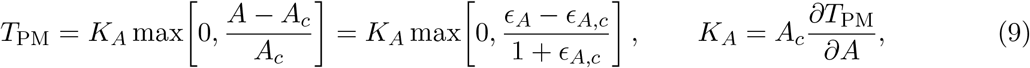

with *K*_*A*_ = 0.25 N*/*m [5]. This relation does not account for channel opening and does not include a rupture criterion. To denote where channels would open and where the PM would rupture, the insets of Figs. 2a and 3a and Supplementary Fig. 2a show horizontal reference levels of 6 and 12 mN*/*m for MscS and MscL activation and 20 mN*/*m as a typical membrane rupture tension.

**Fig. 2.**
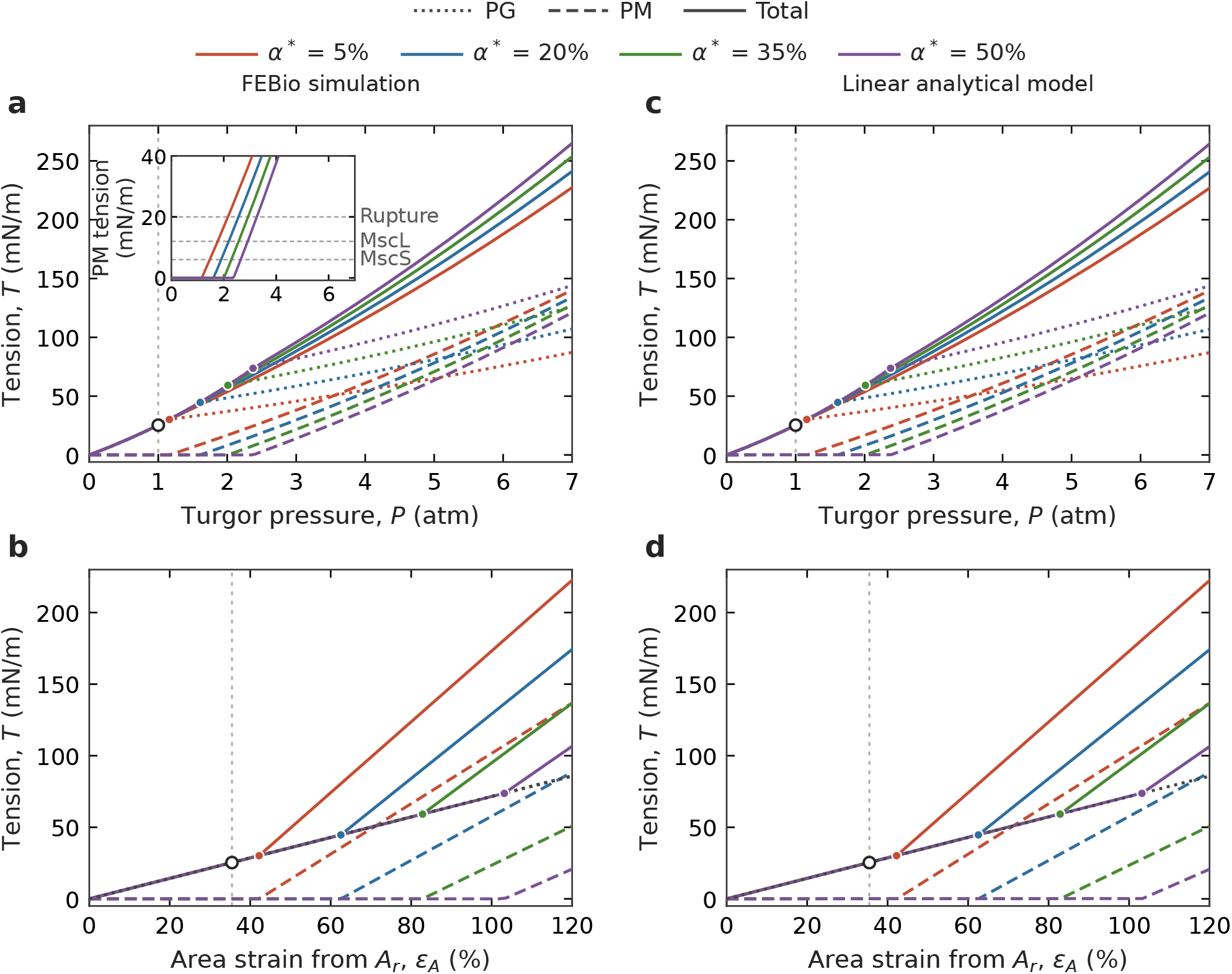
Tension sharing between PG and PM in the linear elastic model. **a**,**b**: FEBio simulations. **c**,**d**: Analytical solution of the same model. **a**,**c**: PG (dotted), PM (dashed), and total wall tension as functions of turgor pressure. **b**,**d**: The same tensions as functions of the area strain *ϵ*_*A*_ = (*A* − *A*_*r*_)*/A*_*r*_ measured from the relaxed PG area *A*_*r*_; the PG curves of all reservoirs coincide and are drawn once (black dotted line). Colors denote the PM excess-area fraction *α*^*^, defined relative to the envelope area *A*_0_ at the 1-atm baseline. The vertical line and open circle mark the baseline (*P*_0_ = 1 atm, *T*_0_ = 25.3 mN*/*m, *ϵ*_*A*,0_ = 0.355); filled circles mark the points of reservoir. depletion. Inset in **a**: FEBio PM tension over the same pressure range, zoomed in on the channel activation range; horizontal lines mark representative MscS (6 mN*/*m) and MscL (12 mN*/*m) activation tension midpoints and the membrane rupture tension (20 mN*/*m). These midpoints describe graded, Boltzmann-type activation rather than sharp thresholdsIncreasing the area reservoir delays the onset of PM tension without changing the PG response before that transition.

**Fig. 3.**
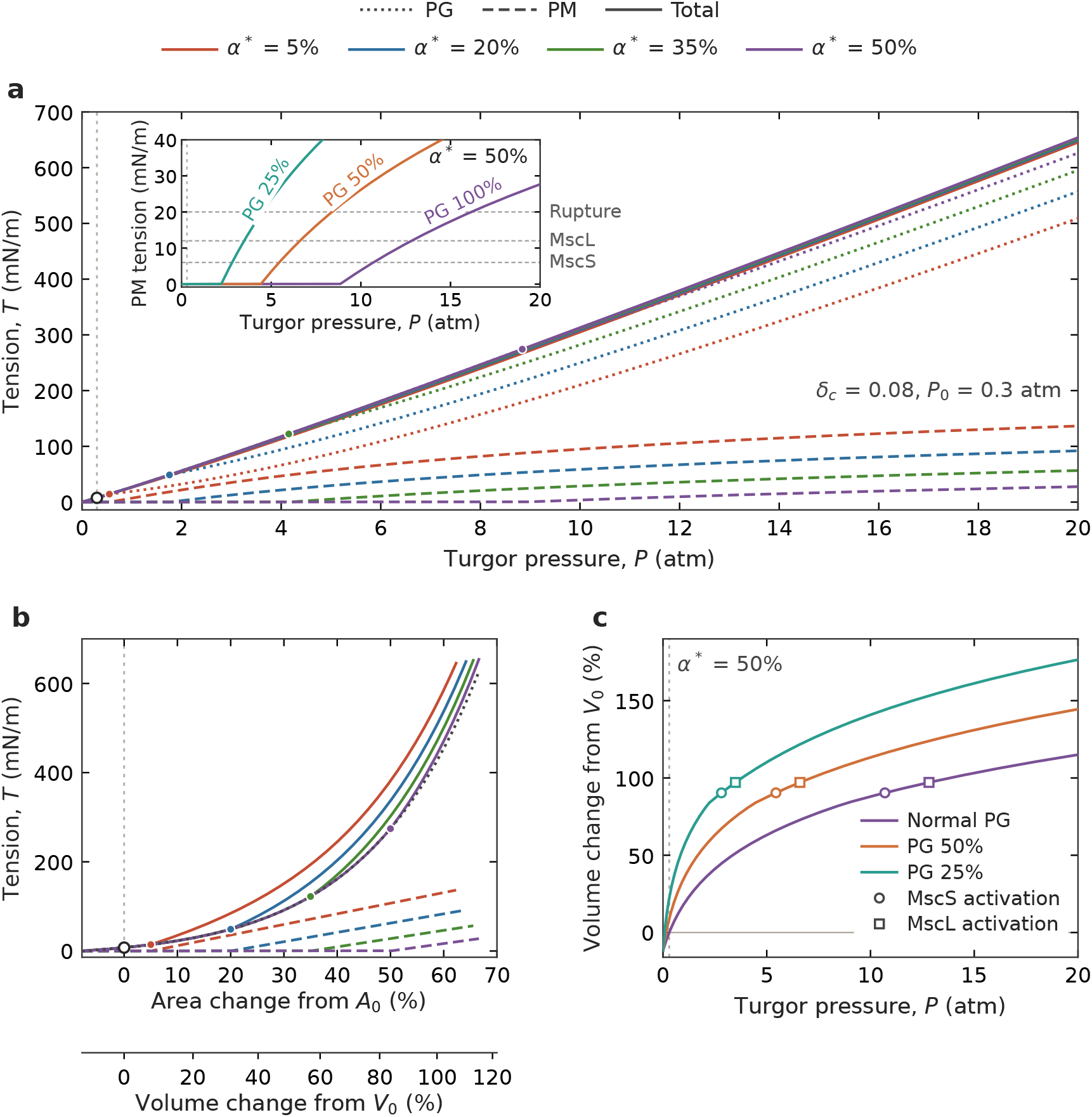
Tension sharing between PG and PM in the model including PG strain hardening and enzymatic PG softening. **a**: PG (dotted), PM (dashed), and total tension for the four reservoirs as functions of turgor pressure. Hardening scale *δ*_*c*_ = 0.08, resting turgor *P*_0_ = 0.3 atm, *r*_0_ = 0.5 µm; the vertical line and open circle mark the resting state, and filled circles mark the points of reservoir. depletion. Inset, PM tension for *α*^*^ = 50% over the same pressure range with intact PG and with PG stiffness reduced to 50% and 25%, as expected after partial digestion by lysozyme or treatment with *β*-lactams. Horizontal lines mark MscS (6 mN*/*m) and MscL (12 mN*/*m) activation tension midpoints and membrane rupture tension (20 mN*/*m). **b**: The same tensions as functions of the envelope area change from the resting area *A*_0_, with the corresponding volume change from the resting volume *V*_0_ on the lower scale, *V/V*_0_ = (*A/A*_0_)^3*/*2^; the PG curves coincide and are drawn once (black dotted line). PG strain hardening increases the PG tension share at large expansion, while the onset of PM tension remains controlled by *α*^*^. **c**, Volume change from *V*_0_ of the untreated cells versus turgor pressure for *α*^*^ = 50% with intact and softened PG; circles and squares mark the representative MscS (6 mN*/*m) and MscL activation tension midpoints, respectively. These midpoints describe graded, Boltzmann-type activation rather than sharp thresholds. Channel opening, solute release, and rupture are not part of the model.

PG softening by sub-lytic concentrations of lysozyme or *β*-lactams was represented by multiplying 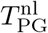 by a factor *s* = 0.5 or 0.25. The relaxed radius *r*_*r*_ and the strain *ϵ*_*A,c*_ at which membrane reservoirs are depleted were kept at the values of the untreated envelope. Areas and volumes are reported relative to the baseline of the untreated cells.

### 2.3 Finite-element simulations

The deformation of the two-layer envelope was numerically simulated using the open-source finite-element-analysis software FEBio version 4.12 [30]. PG and PM were represented as concentric spherical layers with coupled radial displacements, reflecting contact maintained by turgor pressure. A uniform outward pressure was applied to the inner surface and increased quasi-statically from 0 to 7 atm in steps of 0.05 atm. The layers shared the same envelope radius, but their material responses differed: the PG stretched elastically according to Eq. (4), whereas the PM first accommodated expansion by unfolding its excess-area reservoir at zero in-plane tension and then followed Eq. (9). At each pressure step we recorded the mean radius, the area strain of the mesh, and the tensions *T*_PG_ and *T*_PM_; their sum agreed with Eq. (1) to within 0.25%.

### 2.4 Analytical calculations

For each membrane area reservoir size, Eqs. (1)–(9) were evaluated on a dense grid of *ϵ*_*A*_, and the pressure was obtained from the Young–Laplace relation. The baseline strain and the strain at a prescribed pressure were found by bisection; the strains at which PM tension reaches the channel-gating and rupture reference levels follow in closed form from Eq. (9). The analytical model and the plotting scripts are written in Python with NumPy and Matplotlib.

## 3 Results

### 3.1 Tension sharing in a linear elastic envelope

We first treated the PG as a linear elastic layer and the PM as a layer that carries no in-plane tension until its reservoir of membrane area is depleted (Methods). The reservoir size *α*^*^ is the fractional area increase beyond the baseline area *A*_0_ at which the PM becomes taut. Figure 2 shows the FEBio simulations (Fig. 2a,b) next to the analytical solution of the same model (Fig. 2c,d) for *α*^*^ = 5, 20, 35, and 50%, starting from a baseline turgor of *P*_0_ = 1 atm. Geometrically, a membrane reservoir of 50% corresponds to an 84% increase in cell volume when the reservoir is exhausted (Fig. 3b, volume scale).

Below a critical pressure *P*_*c*_, the PM remains slack and the PG carries the full wall tension. As turgor pressure increases, the PG expands elastically while the PM accommodates the same change in envelope geometry by drawing area from its reservoir. During this initial expansion, the PM material itself is not stretched and therefore carries no in-plane tension. Beyond *P*_*c*_, the PM becomes taut, both layers expand together, and the wall tension is shared according to their respective stiffnesses. Because the area modulus of the PM is higher than that of the PG, the tension in the membrane eventually overtakes that of the PG. The composite envelope obeys the Young–Laplace relation, Eq. (1), irrespective of how the tension is distributed between the two layers, and the total tension increases slightly faster than linearly with pressure because the radius of the cell increases with pressure (Fig. 2a,c). When plotted against area strain (Fig. 2b,d), the PM tension kicks in when the envelope area has increased by *α*^*^ relative to *A*_0_; beyond this point both layers are stretched equally, and the total tension rises more steeply with strain.

*P*_*c*_ can be calculated from the model parameters. Combining Eqs. (1), (2), (4), and (8) with *T*_PM_ = 0 gives

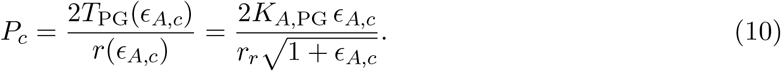

Since *P*_*c*_ marks exhaustion of the reservoir, not channel activation, PM tension has to rise further before it reaches gating thresholds. Larger reservoirs shift *P*_*c*_ to higher pressure: in the linear model *P*_*c*_ = 1.16, 1.61, 2.01, and 2.38 atm for *α*^*^ = 5, 20, 35, and 50%, and PM tension reaches the MscS level (6 mN*/*m) at 1.47, 1.90, 2.29, and 2.65 atm in the FEBio simulations (Fig. 2a, inset).

The FEBio results and the analytical solution differ by less than 0.6 mN*/*m over the full pressure range (Supplementary Fig. 1). This agreement validates the finite-element protocol on a problem with a known solution, and it supports future use of the same protocol for geometries that have no analytical solution, such as rod-shaped *E. coli*, curved *Vibrio* cells, spiral cells, or envelopes with protrusions or invaginations.

### 3.2 Assuming linear PG elasticity underestimates the swelling pressure

Using the PG elastic modulus measured on isolated sacculi [28] in the linear elastic model shows that a 50% membrane reservoir would be exhausted at 2.4 atm turgor pressure, starting from a resting pressure of 1 atm. This is considerably below the pressures expected from the Van’t Hoff relation for the downshifts of 0.4–0.9 Osm applied to cells in stopped-flow experiments [15, 16]. In these experiments, a 50% increase in envelope area (about a twofold increase in volume under the geometric assumptions of ref. [15]) is reached with a 0.6 Osm downshift, which corresponds to about 15 atm at constant cell volume. Cells swell and engage MS channels before osmotic equilibrium is reached, so the pressure at channel activation is lower than 15 atm, but it is clearly higher than 2.4 atm and likely between 5 and 10 atm. A linearly elastic PG layer with the measured modulus is therefore too soft at large extension to explain those results. It is well known however that many biomaterials strain harden [31]. The wall of intact bacterial cells is thus very likely to stiffen under stress ending up being considerably stiffer at resting pressures and above than unpressurized sacculi [29, 32]. We therefore introduced strain hardening of the PG into the analytical model [29] (Methods) and took the resting turgor as *P*_0_ = 0.3 atm, consistent with turgor measured in live *E. coli* [32] and appropriate for cells in LB medium (own unpublished results), in which the stopped-flow measurements were made.

### 3.3 PG strain hardening delays membrane loading

Figure 3a,b shows analytical results obtained using the strain-hardening model with *δ*_*c*_ = 0.08 and *P*_0_ = 0.3 atm. The PG again carries all of the wall tension until the reservoir is exhausted, but strain hardening raises the pressure needed to reach a given envelope area. The PM becomes taut at 0.55, 1.76, 4.15, and 8.84 atm for *α*^*^ = 5, 20, 35, and 50%, and PM tension then rises more slowly with pressure than in the linear model, because the hardening PG takes up a more than linearly growing amount of tension. For the 50% reservoir, PM tension reaches the MscS activation level (6 mN*/*m) at 10.7 atm and the MscL level (12 mN*/*m) at 12.8 atm turgor (Fig. 3a, inset). The hardening scale *δ*_*c*_ is the linear strain over which the PG stiffness increases *e*-fold; *δ*_*c*_ = 0.08 is twice the value inferred within the exponential-hardening model of Albarrán et al. [29]. The onset of PM tension is sensitive to this choice: with *δ*_*c*_ = 0.10 the 50% reservoir is exhausted at 5.5 atm and PM tension reaches the MscS level at 6.5 atm, within the 5–7 atm range estimated above (Supplementary Fig. 2).

### 3.4 Softening the PG lowers the activation pressure

Lysozyme cleaves the glycan strands of the PG and *β*-lactams block its crosslinking. Both treatments reduce PG stiffness. To illustrate the downstream effects of such a treatment, we reduced the PG stiffness to 50% and 25% of its normal value in our model while keeping the relaxed PG area and the amount of PM unchanged (Methods). At the resting pressure the softened PG is stretched further, so part of the reservoir is already used: the resting envelope area increases by 5.9% and 14.3%, respectively. For the 50% reservoir, the PM becomes taut at 4.4 and 2.2 atm instead of 8.8 atm and reaches the MscS level at 5.4 and 2.8 atm instead of 10.7 atm (Fig. 3a, inset). Because PM tension depends only on how far the envelope area exceeds the total relaxed area of the membrane, the PM reaches the MscS and MscL levels at the same cell volume, 90% and 97% above the resting volume *V*_0_ of the untreated cell, for all three PG stiffnesses (Fig. 3c), but softening lowers the turgor at which this volume is reached. In the extreme case, cell expansion due to PG softening may remove the reservoir entirely, in which case the channels would be under tension constitutively and their leakage, rather than turgor and PG elasticity, would set the volume of the cell. We do not model this case.

## 4 Discussion

### 4.1 Implications for mechanosensitive channel function and osmoregulation

Mechanisms of bacterial adaptation to freshwater are critical for the aquatic propagation of both commensal bacteria and pathogens [33–35]. Early views of bacterial osmotolerance held that the PG layer alone was sufficient to restrain cell swelling and prevent lysis [36]. However, identification of two types of tension-activated channels in the *E. coli* cytoplasmic membrane demonstrated that bacteria cannot survive strong osmotic shocks without MscS and MscL [11, 12]. Membrane tension was subsequently viewed as the principal physical factor governing channel activation. Yet it remained unclear when and how membrane tension develops within the multilayered envelopes of Gram-positive and Gram-negative bacteria. Video microscopy showed that a ∼15% linear cell extension under hypotonic shock is corrrelated with osmolyte release through mechanosensitive channels [21]. Faster stopped-flow measurements revealed that an abrupt 600–800 mOsm osmotic downshift produces an approximately twofold increase in cell volume within ∼50 ms, immediately before rapid channel activation [15, 16]. Although the estimated equilibrium turgor pressure under these conditions is 15–20 atm, channels activate well before this level is reached, presumably at roughly half the maximal pressure, rapidly releasing osmolytes and limiting further pressure buildup. If the twofold volume increase corresponds to complete exhaustion of membrane reservoir, one would conclude that a 40–50% excess of inner plasma membrane is under normal conditions maintained as a reservoir within the peptidoglycan [15]. This also implies that, despite non-zero turgor pressure, the plasma membrane does not normally experience lateral tension. Thus, the PG layer shields the PM from tension up to a threshold, at which the membrane reservoir is exhausted and channel-activating tension develops in the PM. The threshold is determined by both the elastic properties of PG and the size of the membrane area reservoir.

Our modeling results suggest that excess PM area plays a central mechanical role in protecting bacteria from damage due to changes in turgor pressure. By accommodating increases in envelope area without immediately accumulating tension, the membrane reservoir prevents channel activation during small turgor fluctuations and avoids unnecessary loss of cytoplasmic contents. Envelope expansion beyond reservoir depletion then rapidly increases PM tension, eventually activating mechanosensitive channels. The size of the PM area reservoir determines when pressure causes membrane tension. Mechanosensitive channels therefore do not directly respond to pressure, but to the membrane tension that develops once excess PM area has been depleted.

The modeling results suggest that mechanosensitive channels may also protect the integrity of the PG. Strain hardening of PG limits envelope expansion and could, in principle, prevent the PM from ever reaching excessive tension. Under extreme loading, however, PG itself is expected to experience failure, leading to wall defects, rupture, and membrane blebbing. Channel opening can release turgor pressure before the PG reaches the damage threshold. The PM area reservoir therefore would serve multiple purposes: it prevents premature channel opening during modest pressure fluctuations while making sure that the membrane will bear tension during severe osmotic shocks. This scenario is consistent with the established view that bacterial mechanosensitive channels are primarily gated by membrane tension to provide emergency protection against rapid hypoosmotic shock [11, 12, 37], but it also solves the puzzle why bacteria can survive up to 50% area increase without membrane rupture. Importantly, the model does not predict a precise physiological pressure threshold for irrecoverable damage; it will depend on the PG constitutive response, channel-gating thresholds, and amount of excess membrane available *in vivo* and on how much loss of cytoplasmic content through the channels the cells can recover from. In a constantly growing bacterium, the PM/PG area ratio will be a dynamic property that is likely to be metabolically regulated in a feedback circuit with cell-wall mechanics to guarantee the proper tension sharing.

Our simulations and calculations describe quasi-static mechanical equilibria: they do not model water influx kinetics, the kinetics of channel opening and solute release, or failure modes of either layer. However, the model clearly predicts a surge of normal turgor pressure right before the onset of lateral tension, a sequence that may have physiological implications. Transient accumulation of released osmolytes in the periplasm could partition the osmotic gradient between the two membranes and buffer the PM [26]. The mechanically coupled PG–OM composite might constitute a mechanically more complex layer than just PG. Future work should include non-spherical geometries, explicit OM mechanics, periplasmic mechanics and transient kinetic effects. Our simplifed model, however, clearly demonstrates the need for a PM reservoir and for precise regulation of PG and PM synthesis in growing bacteria to guarantee resilience to changing environmental conditions. A consequence of the membrane reservoir is the prediction of a substantial surge of turgor pressure before the onset of lateral tension in the PM upon osmotic shock, a sequence of events that may have physiological implications.

## Data and code availability

The FEBio output underlying Fig. 2 and Supplementary Fig. 1, and the code for the analytical model and all figures, are available from the corresponding author upon reasonable request.

## Acknowledgements

This work was supported by the National Science Foundation under grant nos. 2221771 (C.F.S.) and 2221772 (J.D.E.).

## Author contributions

Conceptualization: C.F.S., S.S., J.D.E., O.A., R.G. Methodology: S.S.S.B., O.A., R.G. Software and simulations: S.S.S.B. Formal analysis: S.S.S.B., J.D.E. Funding acquisition: C.F.S., J.D.E., S.S. Supervision: C.F.S., S.S., J.D.E., O.A. Writing of the original draft: S.S.S.B., S.S. Writing, review and editing: C.F.S., S.S., J.D.E., S.S.S.B., O.A., R.G.

## Competing interests

The authors declare no competing interests.

## Supplementary figures

**Supplementary Fig. 1.**
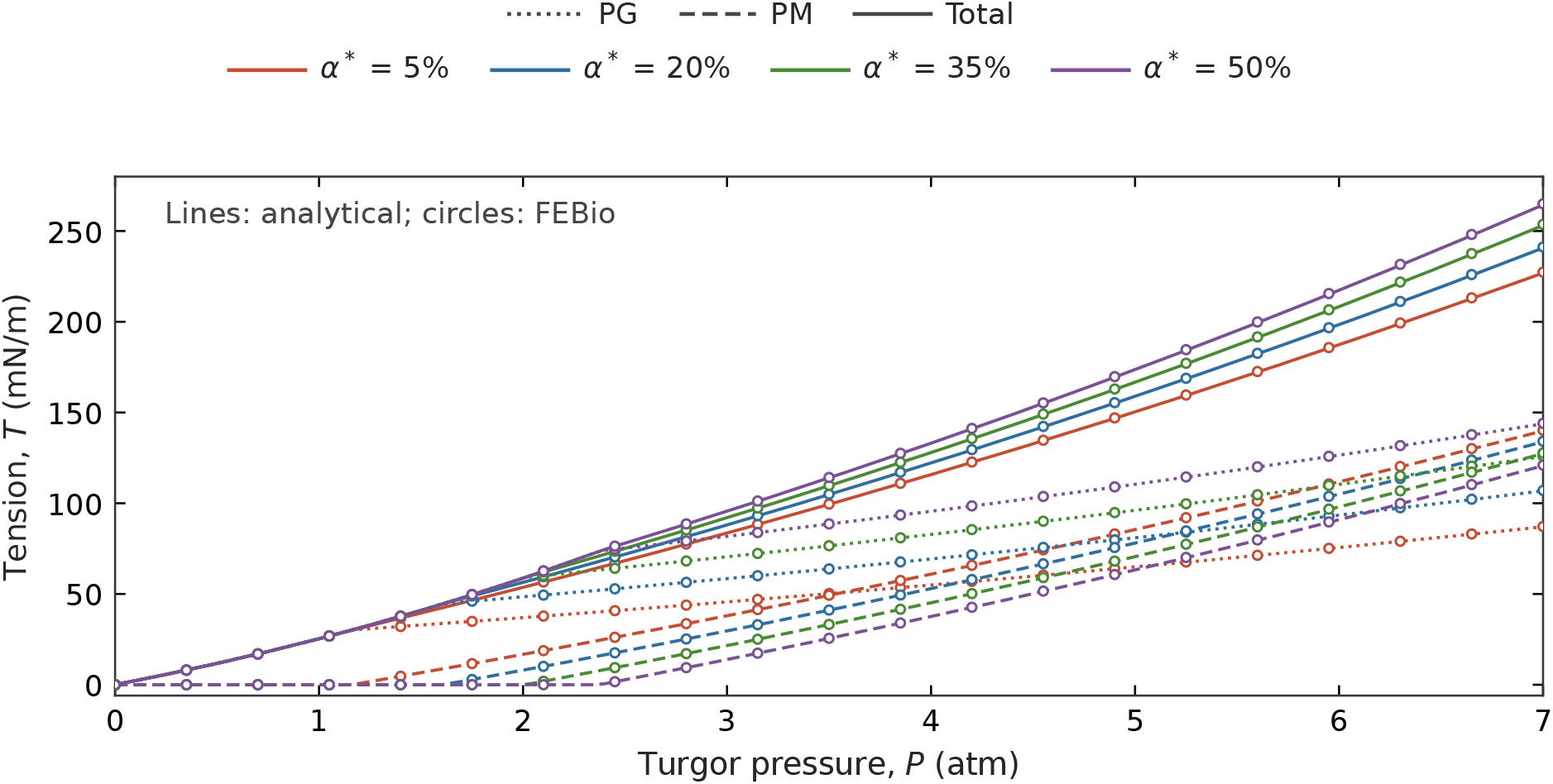
Comparison of FEBio simulations and the analytical calculations of the linear elastic model. PG (dotted), PM (dashed), and total (solid) tension from the analytical model (lines) and from FEBio (circles, every seventh pressure step) for the four reservoir sizes shown in the legend. Baseline *P*_0_ = 1 atm; all parameters as in Fig. 2.

**Supplementary Fig. 2.**
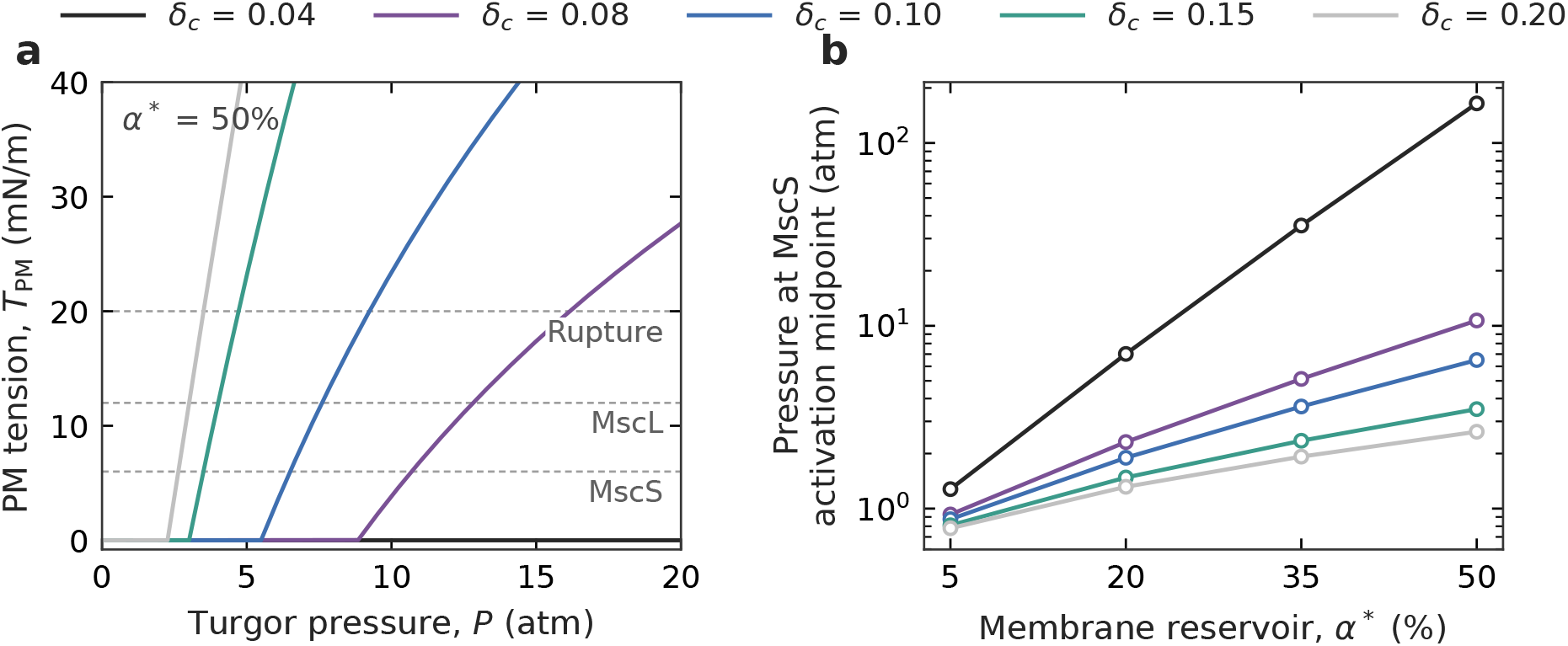
Dependence of PM tension developement on the PG hardening scale *δ*_*c*_. **a**, PM tension for *α*^*^ = 50% at *P*_0_ = 0.3 atm for *δ*_*c*_ between 0.04 and 0.20; *δ*_*c*_ = 0.08 is the value used in Fig. 3. With *δ*_*c*_ = 0.04 the reservoir is not exhausted below 20 atm. **b**, Turgor pressure at which PM tension reaches the MscS activation level (6 mN*/*m) as a function of reservoir size for the same values of *δ*_*c*_ (logarithmic scale).

## Notes

### Competing Interest Statement

The authors have declared no competing interest.

